# c-Rel is required for IL-33-dependent activation of ILC2s

**DOI:** 10.1101/2021.02.14.431188

**Authors:** Aidil Zaini, Thomas S. Fulford, Raelene J. Grumont, Jessica Runting, Grace Rodrigues, Judy Ng, Steve Gerondakis, Colby Zaph, Sebastian Scheer

## Abstract

Group 2 innate lymphoid cells (ILC2s) are emerging as important cellular regulators of homeostatic and disease-associated immune processes. The cytokine interleukin-33 (IL-33) promotes ILC2-dependent inflammation and immunity, with IL-33 having been shown to activate NF-κB in a wide variety of cell types. However, it is currently unclear which NF-κB members play an important role in IL-33-dependent ILC2 biology. Here, we identify the NF-κB family member c-Rel as a critical component of the IL-33-dependent activation of ILC2s. Although c-Rel is dispensable for ILC2 development, it is critical for ILC2 function in the lung, with c-Rel-deficient (*c-Rel*^*–/–*^) mice resistant to papain- and IL-33-induced lung inflammation. We also show that the absence of c-Rel reduces the IL-33-dependent expansion of ILC2 precursors and lower levels of IL-5 and IL-13 cytokine production by mature ILC2s in the lung. Together, these results identify the IL-33-c-Rel axis as a central control point of ILC2 activation and function.

## Introduction

Innate lymphoid cells (ILCs) are important regulators of innate and adaptive immune responses, including inflammatory and allergic responses, as well as in homeostatic processes at barrier tissues (1). ILC subsets are distinguished by distinct developmental pathways, transcription factor expression and production of effector cytokines. Group 1 ILCs (ILC1s) produce IFN-γ and express T-BET, group 2 ILCs (ILC2s) express IL-5, IL-13 and GATA-3, and group 3 ILCs (ILC3s) produce IL-22 and express RORγt (1). Of all ILC subsets, ILC2s are the most important subtype for regulating type 2 immune responses and thus serve key roles in mucosal homeostasis, allergy and anti-helminth immunity. ILC2s primarily reside at mucosal barrier surfaces, highlighting their importance as a first line of defence against invading pathogens. Upon epithelial cell damage, the alarmin IL-33, is released by epithelial cells and binds to a heterodimeric receptor expressed on immune cells, including ILC2s that results in cellular activation (2). Furthermore, IL-33 has been shown to regulate ILC2 mobilisation from the bone marrow to the lungs (3). In addition to IL-33, the epithelial cell-derived cytokines IL-25 and TSLP also have important roles in ILC2 activation and function in response to helminth infections (4), as well as type 2 inflammatory responses such as allergy and asthma (5–7). However, the precise molecular mechanisms of IL-33-induced ILC2 activation remain unclear.

NF-κB comprises a group of transcription factors that play diverse and often critical roles in innate and adaptive immune responses. In mammals, there are five NF-κB family members, namely RelA, RelB, c-Rel, NF-kB1 (p50) and NF-kB2 (p100), all of which share a Rel homology domain (8). NF-κB proteins are comprised of homodimers or heterodimers that normally reside within the cytoplasm in an inactive state. These proteins are rapidly activated in response to diverse upstream signals that typically engage one of two activation pathways: the canonical and non-canonical pathways (9). The canonical pathway activates NF-κB via the IKKβ-dependent phosphorylation and subsequent degradation of the cytoplasmic inhibitibitory protein IκBα, thereby to allow RelA, c-Rel and p50 homo- and heterodimers to translocate to the nucleus and regulate gene expression. In contrast, the non-canonical pathway relies upon IKKα phosphorylation-dependent activation of the NF-kB family members p100 and RelB (10). Although IL-33 has been shown to activate NF-κB in a wide variety of immune cells (2), the precise molecular mechanisms involved are unknown, as are the composition of activated dimers in different cell types, including ILCs.

ST2 is one of the subunits of the IL-33 receptor, with its ligand induced activation triggering MYD88-dependent NF-κB expression, which in turn modulates distinct gene expression programs (2,11,12). Canonical NF-κB signalling has been shown to regulate GATA3 expression in T helper 2 (Th2) cells (13), and T regulatory (Treg) cells (14,15). Furthermore, ST2-IL-33 signalling is associated with the development and function of Th9 cells (16), dendritic cells (17,18), and macrophages (19). IL-33-ST2 signalling is also critical for ILC2 function, although whether NF-κB is required remains unclear (20,21). Recently, non-canonical NF-κB signalling has been shown to be required for IL-33-dependent ILC2s in adipose tissue following death receptor 3 engagement (22), as well as in pulmonary ILC2s upon tumour necrosis factor receptor 2 binding (23). However, adiponectin treatment used to activate the energy sensor AMP-activated protein kinase inhibits the phosphorylation of IKKα/β and IκBα in IL-33-activated adipose-resident ILC2s and impairs IL-13 production (24), suggesting that canonical NF-κB signalling may also play an important role in ILC2 activation and function.

In this present study, we identify a role for the canonical NF-κB family member c-Rel in ILC2 biology under both homeostatic and inflammatory conditions. Our results suggest that while c-Rel is dispensable for ILC2 development in the bone marrow (BM), it is required for peripheral IL-33-dependent ILC2 activation and the development of allergic lung inflammation. These findings point to c-Rel as a potential therapeutic target for treating ILC2-dependent lung inflammation.

## Materials and Methods

### Mice

C57BL/6J mice (wild-type) and c-Rel-deficient mice on C57BL/6J background (*c-Rel*^−/−^) (25) were bred and kept at Monash University. Animals used in this study were 7 to 10 weeks old, with mice maintained under specific-pathogen-free conditions (SPF) conditions. All studies were performed at Monash Biomedicine Discovery Institute (BDI), Monash University in accordance with Monash Animal Ethics Committee (AEC) and Australian National Health and Medical Research Council (NHMRC) guidelines for animal experimentation.

### Electrophoretic mobility shift assay (EMSA)

Nuclear extract preparation was performed as previously described (26). 1-2 μg of nuclear extracts prepared from IL-33-restimulated (6 h) BM-derived ILC2 precursors (ILC2Ps) were incubated with a κB3-specific ^32^P-dATP end-labelled probe, as previously described (27). For supershift analysis, antibodies against p50, c-Rel and RelA were incubated with nuclear extracts on ice for 30 min before adding the radiolabelled probe (27). The samples were incubated for 20 min at room temperature, then 2 μl of gel loading dye Ficoll was added and the samples fractionated on 5% non-denaturing polyacrylamide gels. Gels were dried and exposed to autoradiography.

### Preparation of single cell suspension

BM cells were isolated from femur and tibia by flushing the BM using a 25G needle and passing the cells through a 70-μm strainer to form a single cell suspension. For some experiments, lungs were cut in small pieces and digested in 400 U/ml collagenase IV (Sigma Aldrich), followed by incubation at 37°C for 45 min in complete RPMI media (Life Technologies). The digested lungs were then passed through a 70-μm strainer. Red blood cells (RBCs) in organ cell suspensions were lysed in 1 ml RBC lysis buffer (eBIoscience) for 1 min. After two washes in FACS buffer, the samples were resuspended in a 30% Percoll separation solution and centrifuged at 400 x *g* to enrich the leukocytes, followed by staining for flow cytometry analysis.

### Flow cytometry

Cells were first blocked with purified rat anti-mouse CD16/CD32 (2 μg/ml) (eBioscience) and rat serum (20 μg/ml) (Stem Cell) to prevent non-specific binding of antibodies. Cells were then stained with specific antibodies of interest in the FACS buffer (2% FCS, 1 mM EDTA, and 0.05% azide in PBS). For BM-derived ILC2Ps FACS sorted for culture, the cells were stained in ILC media. Viable cells were identified using the viability dye 7-AAD (eBioscience). The samples were either resuspended in the FACS buffer for acquisition, or fixed overnight at 4°C for intracellular staining the next day using a FOXP3 kit (Tonbo Biosciences). The staining was performed according to the manufacturer’s instructions.

### Cell culture

Bone marrow-derived ILC2Ps or lung ILC2s were sorted by flow cytometry following standard protocols. In short, 5,000 ILC2s were cultured in round bottom plates containing ILC media in the presence of IL-2 (50 ng/ml), IL-7 (10 ng/ml) and IL-25 (100 ng/ml) at d0. Cells were split 1/2 with fresh ILC media plus IL-2 (50 ng/ml), IL-7 (10 ng/ml) and IL-25 (100 ng/ml) at d3 and every second day thereafter over a period of 14 days. On d15, 200,000 cells were plated per well in ILC media in the presence of IL-2 (50 ng/ml) and IL-7 (10 ng/ml) for 24 h. On the following day, the cells were restimulated with IL-33 (10 ng/ml) for 6 h. For some experiments, 5,000 lung-derived ILC2s were cultured overnight in ILC media plus IL-2 (50 ng/ml), IL-7 (10 ng/ml), IL-25 (100 ng/ml) and IL-33 (10 ng/ml) with pentoxifylline (500 ng/ml).

### Lung inflammation model of asthma

Mice were intranasally instilled with 10 μg of papain (Sigma-Aldrich) in 40 μl of PBS under isoflurane anesthesia daily for 3 days. For rIL-33-induced lung inflammation, mice were intranasally injected with 500 ng IL-33 (eBiosciences) daily for 3 days, as previously described (21). 24 h after the last treatment, mice were sacrificed, and their bronchoalveolar lavage (BAL) fluid and lung tissues were collected for flow cytometry analysis and RNA extraction. The left lobe of the lung was obtained and fixed in 10% Formalin solution. The lung tissue was embedded into paraffin blocks and were stained with periodic acid-Schiff (PAS) stain, as previously described (28).

### qPCR

RNA from homogenized tissue was isolated using phenol-chloroform extraction as per standard protocol. RNA from cultured ILC2s was extracted using a NucleoSpin RNA Kit according to manufacturer’s instructions (MACHEREY-NAGEL). The concentration and purity of extracted RNA was measured using a Spectrophotometer NanoDrop 1000 (Thermo Fisher Scientific). 1 μg of RNA was used for cDNA generation using a cDNA conversion kit (Thermo Fisher Scientific). Qpcr was performed using a SYBR green chemistry (Qiagen) on a qPCR system (Rotor-Gene Q Qiagen). Samples were standardized using *Actb*.

### ELISA

ELISA plates were coated with primary antibodies at 1 μg/ml in PBS at 4°C overnight. The plates were washed 4 times using a washing buffer (PBS containing 0.05% Tween 20) (Sigma Aldrich) with 1 min rests between washes. Subsequently, the plates were blocked for 1 h at room temperature with 200 μl PBS containing 10% newborn calf serum (Bovogen). The samples were loaded, followed by a two-hour incubation at room temperature. The plates were washed 5 times, and the secondary antibodies (biotinylated) were loaded (0.5 μg/ml), and incubated for 1 h at room temperature, followed by 8 final washes with washing buffer. TMB substrate (Invitrogen) was added, and the reaction was stopped with HCL. The samples were read at 450 nm using a microplate Epoch spectrophotometer (BioTek) and analyzed using Gen5 2.0 (data analysis software).

### Statistics

All statistical calculations were performed with GraphPad Prism 9 software (GraphPad Software, La Jolla, CA, USA). All graphs represent mean ± SEM. Statistical significance was determined by 2-tailed Student’s t test or one-way-ANOVA. Results were considered statistically significant with p ≤ 0.05.

## Results

### Canonical NF-κB activity in ILC2s

As a start to examining NF-κB involvement in the molecular requirements for IL-33-dependent activation of ILC2s, we examined IL-33-dependent NF-κB activation in ILC2s by electrophoretic mobility shift assay (EMSA). We used flow cytometry to isolate Lin^-^ CD45^+^ c-kit^-^ Sca-1^+^ CD127^+^ CD25^+^ □4β7^+^ ILC2Ps from WT mice and expanded them *in vitro* with IL-2, IL-7, IL-25 and IL-33 for 2 weeks. Expanded ILC2s were rested for 48 hours in IL-2 and IL-7, and subsequently restimulated with IL-33 for 6 h. IL-33 restimulation led to increased nuclear translocation and DNA binding of three distinct NF-κB complexes in ILC2s that are denoted as C1 (lower band), C2 (middle band) and C3 (upper band) (Fig. 1A). Antibody super-shifts revealed that the three complexes were comprised of the canonical NF-κB family members p50 (C1), RelA (C2) and c-Rel (C3) (Fig. 1B). Thus, ILC2s respond to IL-33 stimulation by activating the canonical NF-κB pathway.

**Figure 1.**
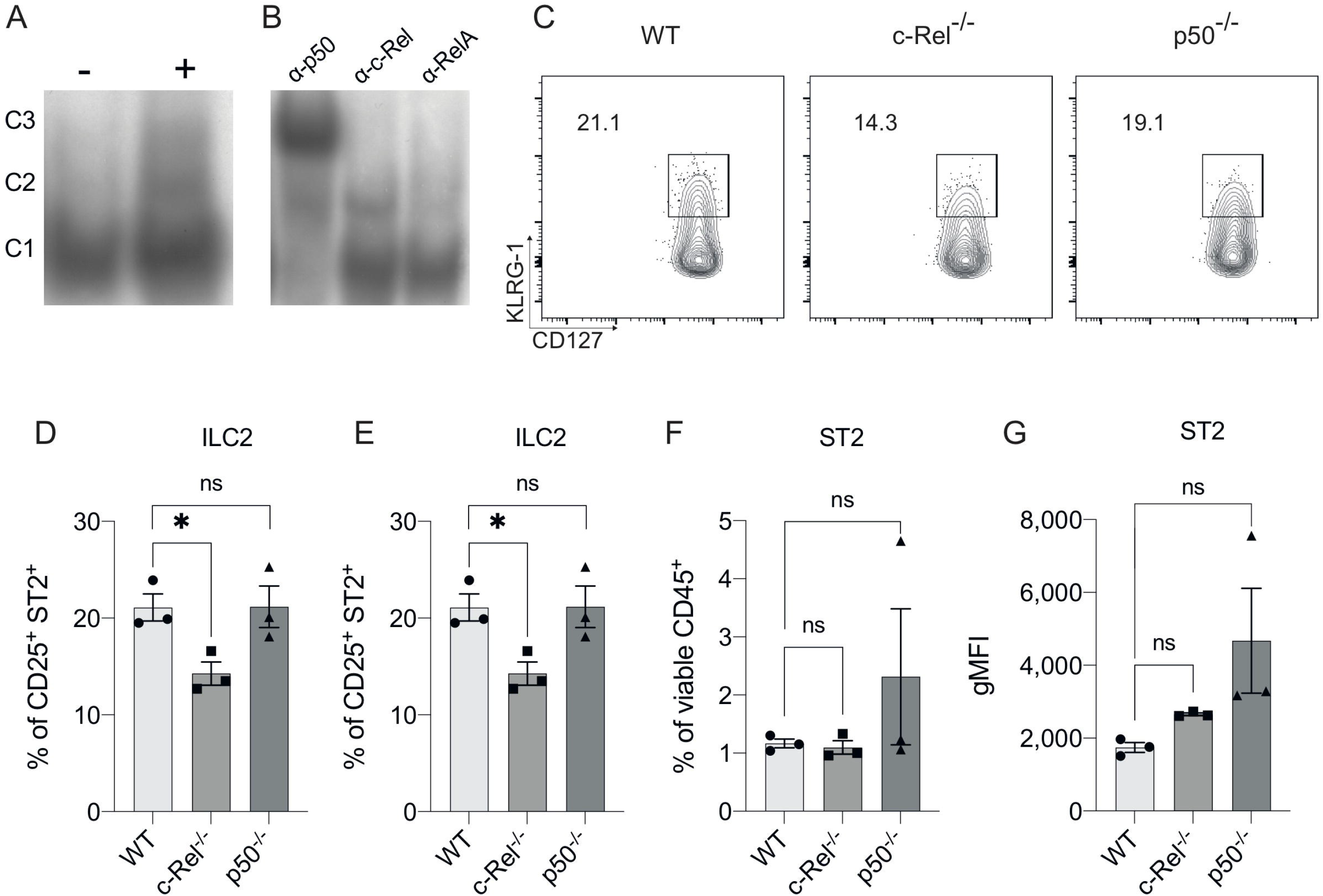
The presence of canonical NF-κB activity in ILC2s. (A) Electrophoretic mobility shift assay (EMSA) analysis in IL-33-stimulated BM-derived ILC2Ps. (B) The specific members of NF-κB were detected via antibody supershift analysis. C1 is composed of p50, C2 is composed of RelA and C3 is composed of c-Rel protein. (C) Representative flow cytometry plot of CD127^+^ KLRG-1^+^ lung ILC2s (gated on viable CD45^+^ Lin^-^ CD90^+^ CD25^+^ ST2^+^ cells) from WT, c-Rel^−/−^ and p50^−/−^ naive mice. (D) Quantification of lung ILC2s among parent CD25^+^ ST2^+^ cells (E) Quantification of lung ILC2s among viable CD45^+^ cells. (F) Quantification of ST2^+^ among viable CD45^+^ cells. (G) Quantification of ST2^+^ cells, expressed as mean fluorescence intensity (MFI). All plots and graphs are representative of at least two independent experiments with 2-3 mice per group. Error bars represent ± SEM. *p ≤ 0.05. ns, non-significant.

We next examined the frequencies of ILC2s in the lungs of c-Rel-deficient (*c-Rel*^*−/−*^) and p50-deficient (p50^*−/−*^) mice. We did not examine the role of RelA as RelA-deficient mice are perinatal lethal (29). We found the frequencies of lung ILC2s were subtly, but significantly reduced by the absence of c-Rel, while p50^*−/−*^ mice had a non-significant reduction in ILC2 frequency (Fig. 1C, D, E). We also found that the absence of c-Rel or p50 had no effect on the expression level or frequency of the IL-33 receptor component ST2 in ILC2s (Fig. 1F, G). Thus, deficiency of the NF-kB family member c-Rel has a minimal yet significant impact on the homeostatic levels of ILC2s in the lungs.

### c-Rel is required for IL-33-dependent ILC2 expansion but not ILC2 development in the bone marrow

We focused our investigation on c-Rel and started by examining the development of immature ILC2s (iILC2s) in the bone marrow of c-Rel-deficient mice during steady state hematopoiesis. Flow cytometric analysis revealed equivalent frequencies of Lin^-^ CD127^+^ α_4_ β_7_ ^+^ CD25^+^ c-Kit^low^ ILC2Ps (30–32) in the BM of control and *c-Rel*^*−/−*^ mice (Fig. 2A, D). Further, cells upstream of ILC progenitors, including common lymphoid progenitors (CLPs), α^4^β^7+^ lymphoid progenitors (αLPs) and common helper innate lymphoid progenitors (ChILPs), were also equally represented in the BM of control and *c-Rel*^*−/−*^ mice (Fig. 2A, B, C). However, we did find that c-Rel was critical for the IL-33-dependent expansion of ILC2Ps. We isolated ILC2Ps from BM of control and *c-Rel*^*−/−*^ mice and stimulated them *in vitro* with IL-2, IL-7 and IL-25. Under this condition, we observed equivalent expansion of control and c-Rel-deficient ILC2Ps (Fig. 2E), although to a significantly lesser extent than when IL-33 was present. However, unlike WT ILC2Ps, c-Rel-deficient ILC2Ps failed to expand in response to IL-33. Thus, c-Rel is critical for the IL-33-responsiveness of ILC2Ps.

**Figure 2.**
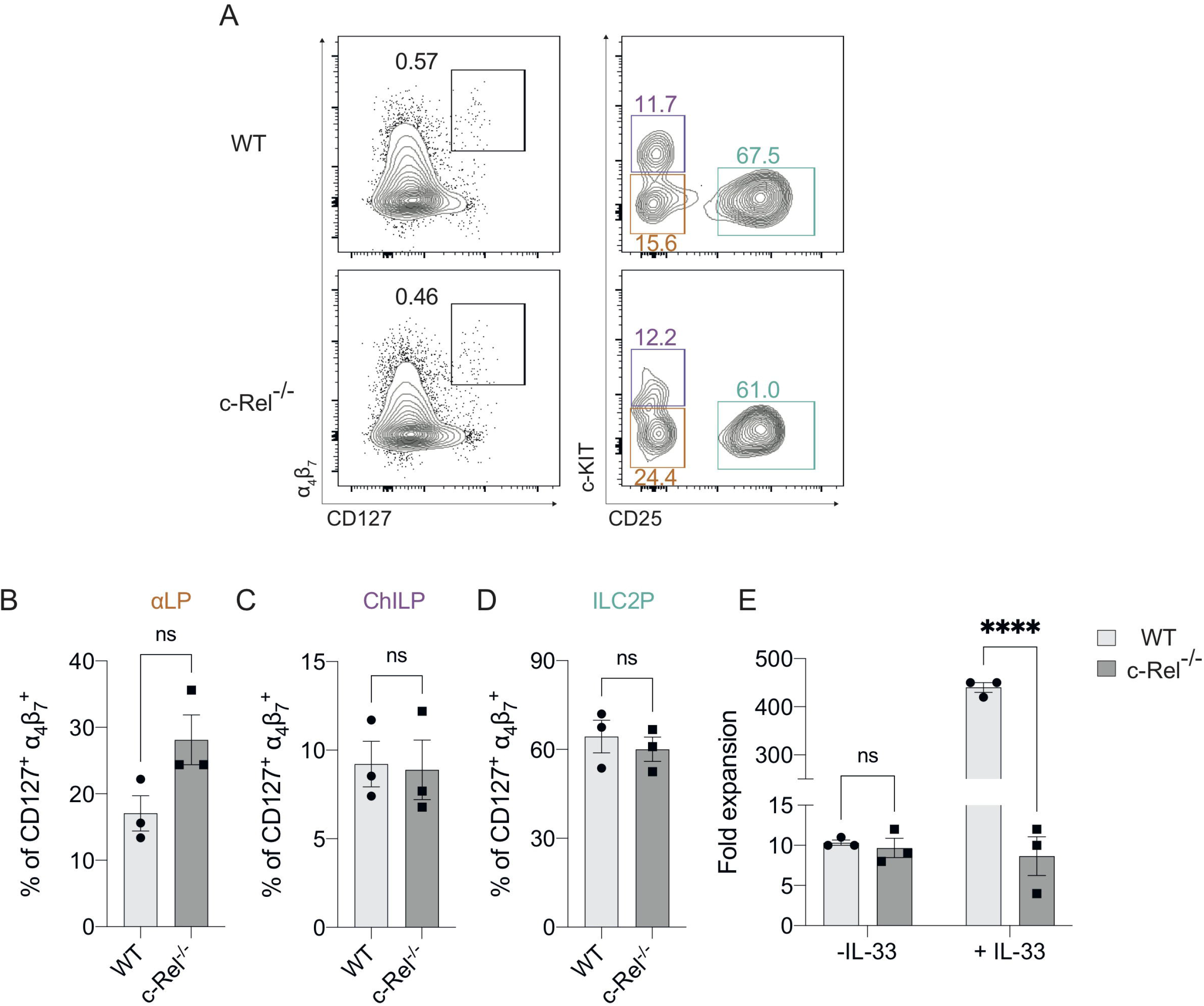
The development of BM-derived immature ILC2s are independent of but require c-Rel for their IL-33-dependent expansion. (A) Representative flow cytometry plot of α4β7^+^ CD127^+^ immature BM ILC2s (left) (gated on viable CD45^+^ Lin^-^ cells) and specific ILC2 progenitors (gated on parent α4β7^+^ CD127^+^ cells). α-lymphoid progenitors (αLPs) were defined as c-Kit^low^ CD25^-^, whereas common helper innate lymphoid progenitors (ChILPs) and ILC2 precursors (ILC2Ps) were c-Kit^high^ CD25^-^ and c-Kit^low^ CD25^+^, respectively. (B, C, D) Quantification of αLP, ChILP and ILC2P among α4β7^+^ CD127^+^ cells, respectively. (E) Quantification of fold expansion derived from cell count on ex vivo BM-derived ILCP cultures. FACS-sorted BM ILC2Ps were cultured in ILC media in the presence of IL-2, IL-7 and IL-25 for 14 days. On d15, expanded cells were cultured in only IL-2 and IL-7 overnight. On d16, expanded cells were re-stimulated with or without IL-33 (control) for 6 h, followed by cell count. All plots and graphs are representative of at least two independent experiments with three mice per group. Error bars represent ± SEM. *p ≤ 0.05, **p ≤ 0.01, ****p ≤ 0.01. ns, non-significant.

### c-Rel regulates papain-induced lung inflammation

IL-33 is maintained in the nucleus of epithelial cells under homeostatic conditions, but is released following tissue damage (2). Therefore, we used a model of allergic lung inflammation to determine if c-Rel was required for ILC2 function under inflammatory conditions. We treated control WT and *c-Rel*^*−/−*^ mice intranasally with the protease allergen papain on days 0, 1 and 2, and examined the acute, ILC2-dependent inflammatory response in the lungs on day 3 (32–34). Strikingly, *c-Rel*^*−/−*^ mice showed reduced papain-induced lung inflammation, with a significantly diminished influx of lymphocytes into the bronchoalveolar lavage (BAL) and significantly fewer eosinophils (Fig. 3A, B, C). Like homeostatic conditions in the BM, we did not find any significant difference in the frequencies of lung ILC2s in *c-Rel*^*−/−*^ and control mice (Fig. 3D). However, we observed significantly reduced *Il5* and *Il13* expression in the absence of c-Rel, suggesting that c-Rel plays an important part in the expression of these type 2 inflammatory proteins. (Fig. 3E, F). As damaged epithelial cells also produce IL-25 and IL-33 in response to injury, which could in turn activate ILC2s in a paracrine manner (2) to produce IL-5 and IL-13, we examined their involvement in this model. Importantly, we found that expression levels of *Il25 and Il33* were equivalent between *c-Rel*^*−/−*^ mice and control mice (Fig. 3G, H), showing that c-Rel does not regulate *Il25* and *Il33* expression in epithelial cells in our experiments. Collectively, these data suggest that c-Rel is an important regulator of ILC2-dependent lung inflammation.

**Figure 3.**
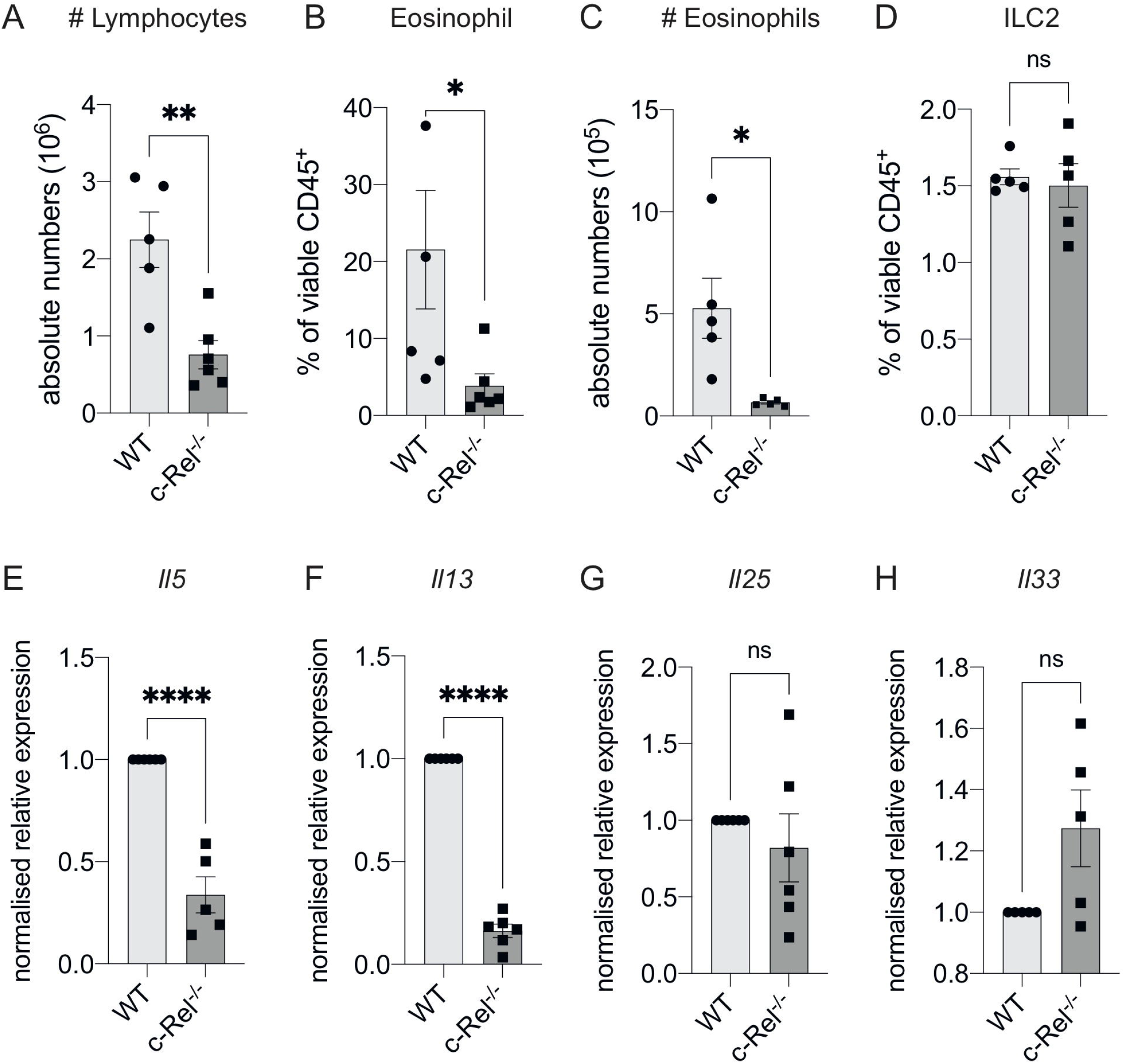
c-Rel deficiency reduces papain-induced lung inflammation. (A) Absolute numbers of CD45^+^ lymphocytes in the bronchoalveolar lavage (BAL) fluid following papain challenge for three days. (B) Quantification of Siglec-F^+^ CD11c^-^ eosinophils in BAL among viable CD45^+^ cells. (C) Absolute numbers of BAL eosinophils. (D) Quantification of CD127^+^ KLRG-1^+^ lung ILC2s (gated on Lin^-^ CD90^+^ CD25^+^ ST2^+^) among viable CD45^+^ cells. (E, F, G, H) qPCR analysis of the indicated genes from lung tissue following papain challenge. All graphs are representative of at least two independent experiments with 5-6 mice per group. Error bars represent ± SEM. *p ≤ 0.05, **p ≤ 0.01 ns, non-significant.

### c-Rel is indispensable for IL-33-dependent lung inflammation and activation

As papain treatment induces expression of the ILC2-inducing cytokine IL-33, and given that IL-33 treatment of ILC2s leads to c-Rel activation, we next tested whether c-Rel was required for the direct activation of ILC2s by IL-33 *in vivo*. We treated control WT and *c-Rel*^*−/−*^ mice intranasally with rIL-33 on day 0, 1 and 2, and analyzed lung responses on day 3. Consistent with our lung inflammation model using papain, *c-Rel*^*−/−*^ mice displayed reduced IL-33-dependent inflammation in the lungs, with reduced infiltration of lymphocytes and eosinophils into the BAL (Fig. 4A, B, C), reduced frequencies of ILC2s in the lungs (Fig. 4D), and lower expression levels of *Il5* and *Il13*, but not *Il25*, in lung tissue (Fig. 4E, F, G). IL-33-induced goblet cell hyperplasia and mucus production in the lungs lacking c-Rel was also severely reduced (Fig. 4G). Further, *ex vivo* activation of lung-derived ILC2s with IL-2, IL-7, IL-25 and IL-33, resulted in the c-Rel-dependent production of IL-5 and IL-13 (Fig 4H, I). Additionally, in the presence of pentoxifylline, an NF-κB inhibitor with selectivity for c-Rel over other family members (35,36), c-Rel-sufficient ILC2s failed to produce IL-5 and IL-13 (Fig 4H, I), in a similar manner to c-Rel-deficient ILC2s. Together, these observations suggest that c-Rel is critical for IL-33-dependent activation of ILC2s during lung inflammation.

**Figure 4.**
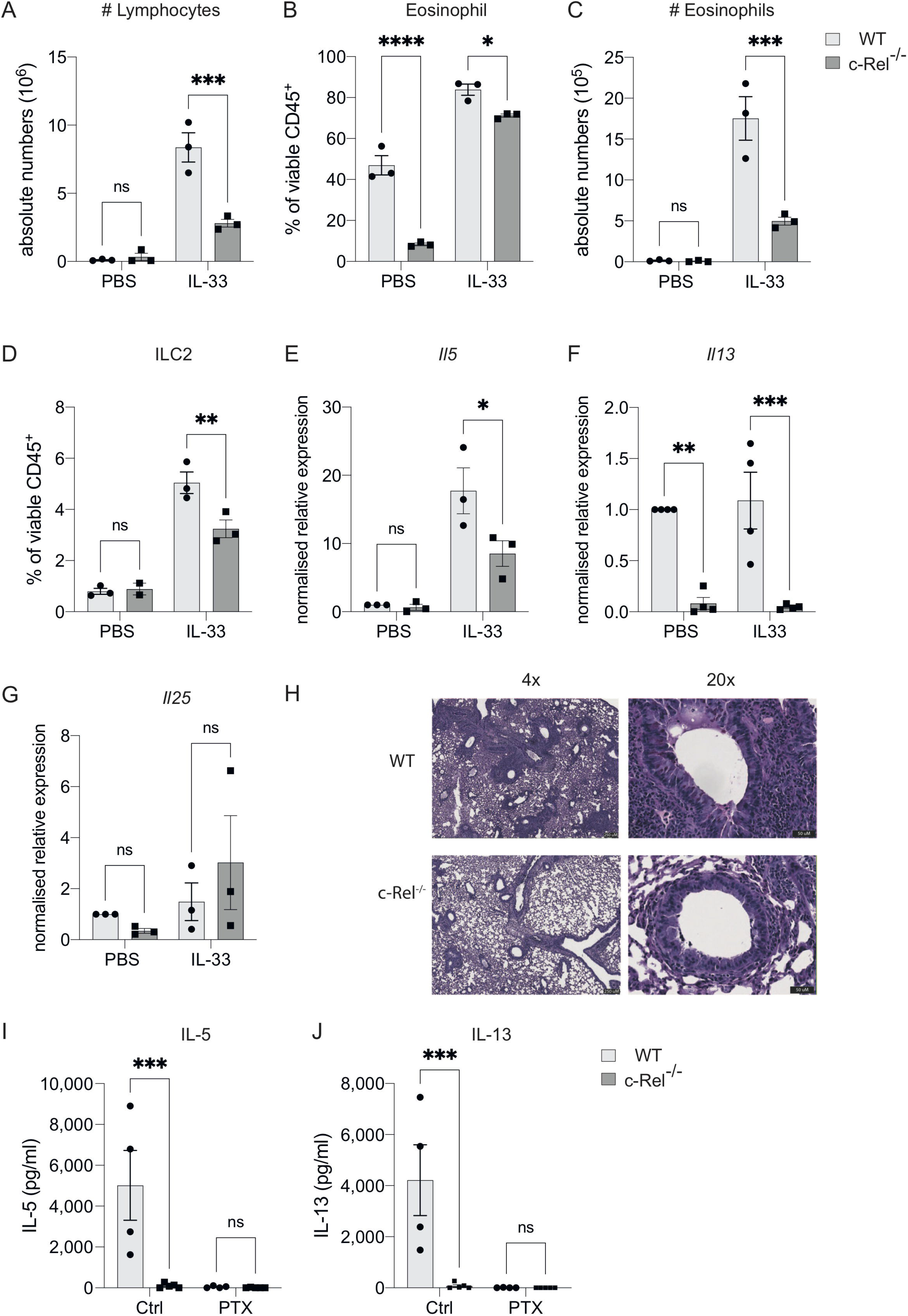
c-Rel is critical for ILC2-mediated lung inflammation, induced by IL-33. (A) Absolute numbers of CD45^+^ lymphocytes in the bronchoalveolar lavage (BAL) fluid following PBS (control) or rIL-33 administration for three days. (B) Quantification of Siglec-F^+^ CD11c^-^ eosinophils in BAL among viable CD45^+^ cells. (C) Absolute numbers of BAL eosinophils. (D) Quantification of CD127^+^ KLRG-1^+^ lung ILC2s (gated on Lin^-^ CD90^+^ CD25^+^ ST2^+^) among viable CD45^+^ cells. (E, F, G) qPCR analysis of the indicated genes from lung tissue following PBS or rIL-33 administration. (H) PAS-stained lung histology visualised using 4x (left) and 20x (right) magnification. (I, J) ELISA analysis of the indicated cytokine from the supernatant of ex vivo lung ILC2s cultures in the presence of IL-2, IL-7, IL-25 and IL-33, with or without a c-Rel inhibitor, pentoxifylline (control). All graphs are representative of at least two independent experiments with 3-5 per group. Error bars represent ± SEM. *p ≤ 0.05, **p ≤ 0.01, ***p ≤ 0.001 ns, non-significant.

## Discussion

Although IL-33 has been shown to be a potent activator of ILC2s during inflammatory responses, the molecular mechanisms responsible for its regulation are poorly understood. While the primary source of IL-33 is epithelial cells (2), other cell types such as NKT cells and alveolar macrophages (37) have also been shown to produce IL-33. Previous studies have established a role for several factors for a number of transcription factors such as GFI1 (38), ETS1 (39), TCF1 (40,41), G9a (32), and non-canonical NF-κB factors (22,42) as important regulators of ILC2 biology; however, the roles of canonical NF-κB proteins have not been directly examined in ILC2s. The present study identified the presence of canonical NF-κB family members p50, RelA and c-Rel activity in ILC2s in following IL-33 stimulation. We found that while c-Rel is dispensable for early ILC2 precursor development in BM, it is required for optimal ILC2P expansion and proliferation. Furthermore, our results suggest that c-Rel regulates ILC2 activation and function in response to papain- or rIL-33-induced lung inflammation. Overall, the present study identifies a role for c-Rel as a regulator of IL-33-dependent ILC2 function, particularly during lung inflammatory responses.

We observed an increase in nuclear localisation and DNA binding of each canonical NF-κB family member p50, RelA and c-Rel in ILC2s upon IL-33 stimulation. Typically, NF-κB can act as a homodimer (e.g. c-Rel:c-Rel), and/or a heterodimer (e.g. c-Rel:p50) (2,9). Thus, further characterization of the functional roles in different combinations of NF-κB family members in IL-33-stimulated ILC2s is needed. In response to IL-33, the expression of canonical NF-κB in our murine activated ILC2s is consistent with previous findings for human IL-1β-primed ILC2s that require IKK-mediated activation of NF-κB (43). In addition to canonical NF-κB signals, non-canonical NIK-dependent NF-κB signalling involving p100 and RelB activation occurs in pulmonary ILC2s, specifically in response to alveolar macrophage-derived TNF-α via TNFR2 (42). Notably, the expression of the canonical NF-κB genes in TNF-α-stimulated ILC2s is either unchanged or reduced (42). Thus, we speculate that the activation of canonical NF-κB signals in pulmonary ILC2s is largely dependent on the pathway downstream of IL-33 mediated by c-Rel, whereas the non-canonical signals are activated in response to other activating cytokines such as TNF-α.

Unlike conventional cytokines, IL-33 is typically released by epithelial cells following damage to barrier tissue and binds to ST2 that forms a heterodimer with IL-1RAcP (44,45) to serve as an alarmin in response to inflammation and infection (46–48). During the steady state, IL-33 is constitutively expressed in both human (49) and mouse airway epithelial cells (50). Our results show that in steady-state conditions, the absence of either c-Rel or p50 does not impair the proportion of ILC2s expressing ST2. However, we observed that the homeostatic development of pulmonary naive ILC2s is somewhat compromised when c-Rel is absent. This indicates that c-Rel-dependent immature ILC2s in the lungs develop independently of ST2 expression. Alternatively, c-Rel may be a prerequisite for IL-33-mediated egress of ILC2s from BM to the lungs (3), which can potentially explain why the accumulation of naive lung ILC2s is somewhat reduced in the absence of c-Rel. Notably, there is only a minimal reduction in ILC2s in the lungs of naive mice lacking c-Rel, suggesting that c-Rel is not absolutely required for normal naive pulmonary ILC2 development. This is in line with our earlier observation, in which minimal NF-κB activity is detected in non-activated ILC2s in the absence of IL-33, but its activation is increased upon IL-33 stimulation. Such an observation is consistent with prior studies that have demonstrated the expression and nuclear activity of NF-κB in resting, naive T cells is minimal (51–53).

ILC development occurs in the BM and immature ILCs then migrate to specific tissues (54). A previous study showed that IL-33/ST2 signalling is not required for ILC2 development, but leads to fewer peripheral ILC2s due to reduced egress from the BM (3). Our characterization of αLP, ChiLP and ILC2P cell populations also revealed that c-Rel is not absolutely required for early development of BM ILC2 progenitors during hematopoiesis. Similar to IL-33- and ST2-deficient mice, we also observed a slight reduction in immature ILC2s in the lungs of c-Rel−/− mice, suggesting that some aspects of BM egress or homeostasis are also regulated by c-Rel. However, we demonstrated that *ex vivo* BM-derived ILC2Ps lacking c-Rel are severely defective in their ability to expand in response to IL-33, highlighting an intrinsic role of c-Rel as part of IL-33 downstream activation pathway in ILC2s. Thus, c-Rel appears to primarily regulate IL-33-dependent ILC2 activation.

Numerous studies have demonstrated that both human (55–57) and mouse ILC2s are intimately associated with respiratory and allergic diseases, including asthma and that activation of ILC2s requires IL-33 signals (58,59). Expression of *Il33* or *Il1rl1* genes also correlates with susceptibility to asthma (19,60–62), as well as the type 2 response in murine experimental asthma models (2,58,59), further supporting a crucial role of IL-33-mediated ILC2s activation during allergy. As the loss of c-Rel impairs *ex vivo* IL-33-dependent ILC2 proliferation and expansion, we questioned whether c-Rel could also be responsible for ILC2-mediated allergic lung responses. The papain-induced lung inflammation model has been shown to induce ILC2-dependent IL-5 and IL-13 production and airway eosinophilia (32–34). We found that c-Rel is critical for the development of papain-induced allergic lung inflammation. We observed a significantly reduced airway eosinophilic response and lower levels of *Il5* and *Il13* expression in the lungs of c-Rel-deficient mice. Surprisingly, the frequencies of lung ILC2s lacking c-Rel remain at the equivalent levels seen in control mice, implying that c-Rel is critically required for ILC2 activation in producing IL-5 and IL-13, but not ILC2 development or survival in response to papain.

Blockade of IL-33 signalling has been shown to limit the development of ILC2-mediated chronic asthma (63), with the administration of IL-33 activating ILC2-dependent lung inflammatory responses and goblet cell hyperplasia (12). Similar to our results with papain, intranasal treatment of c-Rel-deficient mice with rIL-33 also resulted in reduced eosinophilia and lower expression of *Il5* and *Il13* in the lungs. In contrast to the papain results, we observed a reduction of lung ILC2s in c-Rel-deficient mice responding to rIL-33. This might be due to the direct effects of rIL-33 on lung ILC2s that could provide a stronger stimulation than the papain allergen. Collectively, it appears that c-Rel is important for ILC2-dependent lung inflammation in response to papain and IL-33, with this finding confirming and extending previously published work on the role of c-Rel in promoting ovalbumin-alum induced airway inflammation (64).

c-Rel is responsible for many important immune cell functions controlled by regulating the expressions of a wide range of genes (65), including *Il13* (66) and is therefore also likely to be implicated in IL-13-producing Th2-mediated responses associated with asthma and allergy. Further, the loss of p50 alone, or in combination with the absence of c-Rel is associated with impaired effector CD4 T cell survival (67) and Th2 cell functions following ovalbumin challenge (13), respectively. p50-deficient mice failed to mount lung allergic responses due to reduced GATA3 expression in Th2 cells and impaired type 2 cytokine secretion (13). Given p50 can dimerize with c-Rel (65), this points to a potential role of c-Rel in Th2 cell functions, including the possible involvement of impaired c-Rel dependent Th2 cells functions as part of the explanation for the defective inflammatory responses observed in our c-Rel-deficient mice. Although we showed that the deletion of c-Rel protects the mice from papain- and rIL-33-induced lung inflammation, at this time we cannot completely rule out whether c-Rel-intrinsic roles in Th2 cells are also key mediators of the responses we observed, and this remains to be further explored through the use of c-Rel ablation on a Rag knockout background. However, a comparative analysis of IL-5 and IL-13 production in murine experimental asthma showed that major producers of these type 2 cytokines are ILC2s, rather than Th2 cells (5). Furthermore, in line with our data showing that in cultures of *ex vivo* c-Rel-deficient mice ILC2s or pulmonary WT ILC2s treated with the c-Rel inhibitor, pentoxifylline, both sets of cultured cells failed to produce IL-5 and IL-13. This makes it very likely that the response observed in our *in vivo* asthma model experiments is largely mediated by c-Rel-dependent ILC2s.

NF-κB is not only expressed by immune cells, but also non-immune cells such as epithelial cells. It has been previously shown that transgenic mice expressing active IκB kinase (IKK) β in airway epithelial cells develop allergic airway disease due to activation of NF-kB signalling that in turn elevates Th2 and ILC2 responses during lung inflammation (68). Conversely, selectively preventing IKK-dependent NF-κB activation in mouse intestinal epithelial cells impairs Th2 responses following helminth infection, resulting in susceptibility (69). Thus, these findings point to an important role of epithelial cell-intrinsic NF-κB signals in mediating the type 2 response. Since we used global c-Rel knockout mice in the present study, we acknowledge that the immune response generated in our lung inflammation models may not be entirely ILC2-intrinsic if the c-Rel-mediated epithelial cell activation is contributing to the response observed. However, we observed equivalent expression of the epithelial cell-derived cytokines *Il25 and Il33* in papain-induced inflamed lungs of control and *c-Rel*^*−/−*^ mice, indicating that c-Rel deletion in airway epithelial cells does not affect upstream activation component of the inflammatory response.

Asthma and allergic diseases are rapidly increasing worldwide, highlighting a crucial need for a novel anti-inflammatory drug that can efficiently modulate the type 2 response, known to be central to immune responses responsible for lung inflammation. In addition to Th2 cells, ILC2s are key producers of IL-5 and IL-13. Unlike Th2 cells, ILC2s mainly reside at the interface between the host and environment, such as in the submucosa of lungs, initiating inflammation in response to allergens. Accordingly, the localization of ILC2s provides a strategic approach to specifically target ILC2s to attenuate airway hyperreactivity associated with asthma. Our results suggest that a c-Rel-specific inhibitor such as the IT-603 and IT-901 compound (70,71) may offer a novel therapeutic strategy to inhibit IL-33-induced ILC2 activity in diseases such as asthma and allergies. In conclusion, our results demonstrate that c-Rel is an important regulator of IL-33-induced ILC2s during allergic lung inflammation.

## Author contribution

Designed the study and conceptualisation: CZ, SS, and AZ. Performed the experiments: AZ, SS, JR, GR, and JN. Analyzed and interpreted experimental data: AZ, and SS. Performed the EMSA experiment: RJG. Provided mouse strains: TSF, and SG. Wrote and drafted the manuscript: AZ. Provided comments, edited and reviewed the manuscript: CZ, SS, SG and AZ. The authors declare that no conflicts of interest exist.

## Acknowledgments

We would like to thank the Monash animal facility, the Monash Flow Core facility and the Monash Histology Platform facility for their excellent technical support and assistance.

---

**Table 1.** Flow cytometry antibodies

| Antibody | Clone | Supplier |
| --- | --- | --- |
| CD3ε | 145-2C11 | Tonbo Biosciences |
| CD4 | GK1.5 | Tonbo Biosciences |
| CD5 | 53-7.3 | Tonbo Biosciences |
| CD8 | 53-6.7 | Tonbo Biosciences |
| CD11b | M1/170 | Tonbo Biosciences |
| CD11c | N418 | eBioscience |
| CD19 | eBio1D3 | eBioscience |
| CD45R (B220) | RA3-B62 | eBioscience |
| MHCII | M5/114 | eBioscience |
| Ly6G/C | RB6-8C5 | eBioscience |
| Ter119 | TER-119 | Tonbo Biosciences |
| NK1.1 | PK136 | eBioscience |
| α<sub>4</sub>β<sub>7</sub> | DATK32 | eBioscience |
| Sca-1 | D7 | eBioscience |
| CD25 | P661.5 | eBioscience |
| c-Kit (CD117) | 2B8 | eBioscience |
| CD127 | SB/119 | BD Bioscience |
| CD45 | 30-F11 | eBioscience |
| ST2 (IL-33R) | RMST2-2 | eBioscience |
| KLRG1 | 2F1 | eBioscience |
| CD25 | P661.5 | eBioscience |
| CD90.2 | 53-2.1 | eBioscience |
| CD127 | SB/119 | BD Bioscience |
| CD45 | 30-F11 | eBioscience |
| Siglec-F | E50-2240 | BD Bioscience |

**Table 2.**
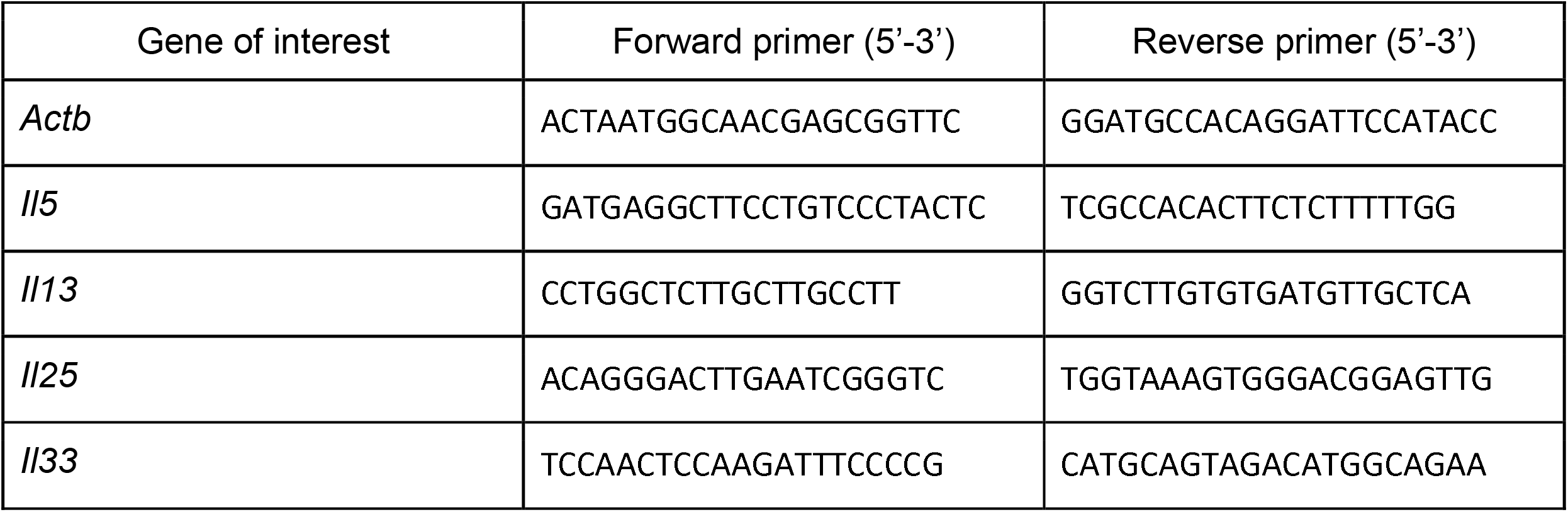
qPCR primers

